## Supplementary figures and legends for "Filamentous recombinant human Tau activates primary astrocytes via an integrin receptor complex"

**Supplementary information**

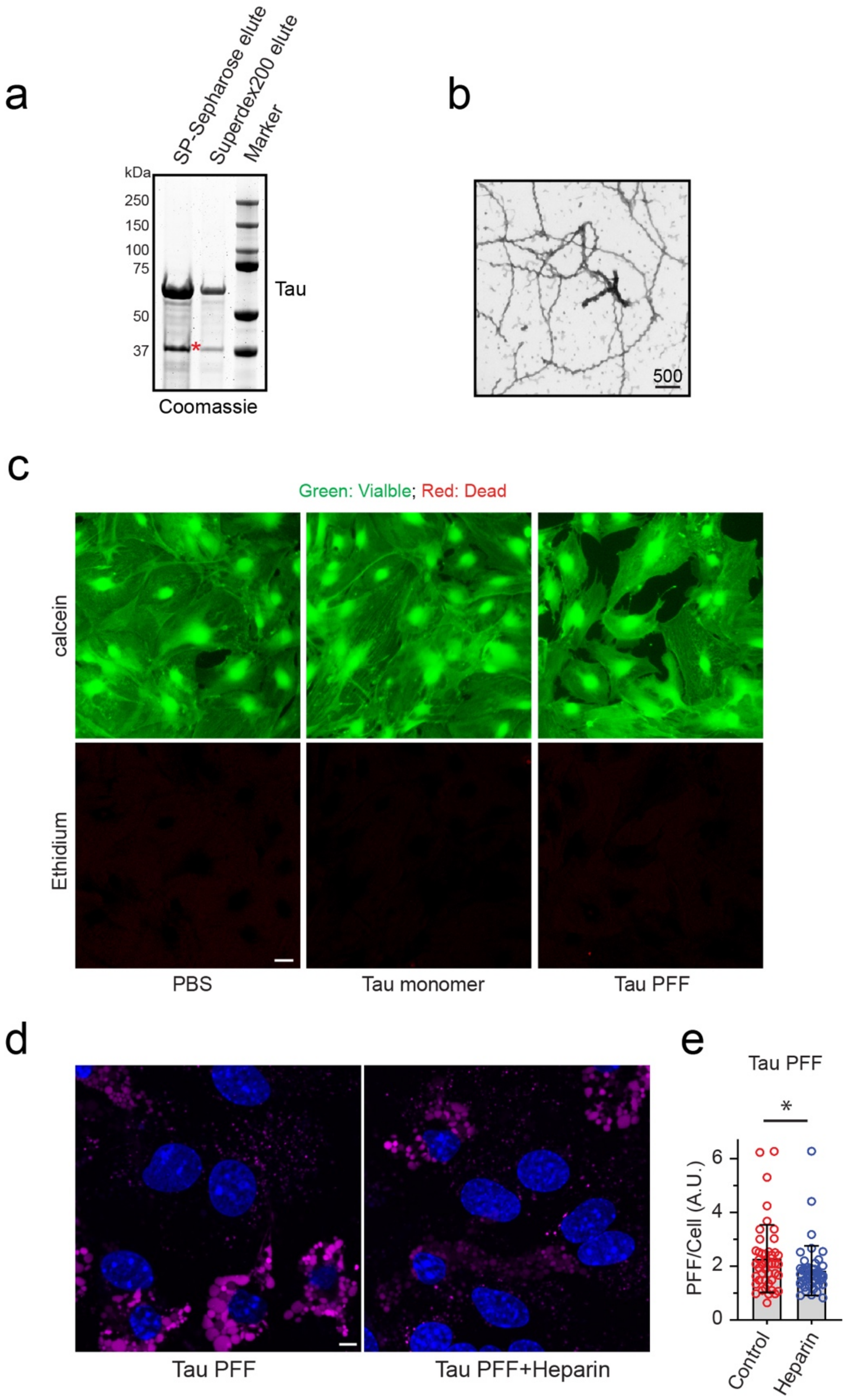


**Figure S1. Tau PFF endocytosis by primary astrocytes is independent of HSPG.**

(**a**) SDS-PAGE and Coomassie blue staining show the purified untagged Tau 2N4R protein. Protein eluates from SP-Sepharose and Superdex 200 were analyzed. Asterisk indicates a degradation product.

(**b**) A representative negative stain EM picture of the assembled Tau PFF before sonication. Scale bar, 500nm.

(**c**) Short-term treatment of astrocytes with Tau monomer or PFFs did not induce cell death. Primary astrocytes treated with PBS, Tau monomer or PFF (200 nM) for 6 h were stained with a green-fluorescent dye calcein-AM to label viable cells and a red-fluorescent dye ethidium homodimer-1 to label dead cells. Scale bar, 10 μm.

(**d**) Astrocytic uptake of Tau PFF is not significantly inhibited by heparin. Astrocytes were incubated with Alexa 594-conjugated Tau PFF (Magenta) for 2 h in the absence or presence of 50 μg/ml heparin. Cells were stained with Hoechst (Blue) before confocal imaging.

(**e**) Quantification of Tau PFF fluorescence signal in individual cells in experiments shown in **d**. Mean ±SEM, n=3 biologically independent experiments. *, $p<0.05$ by two-tailed unpaired student t-test.

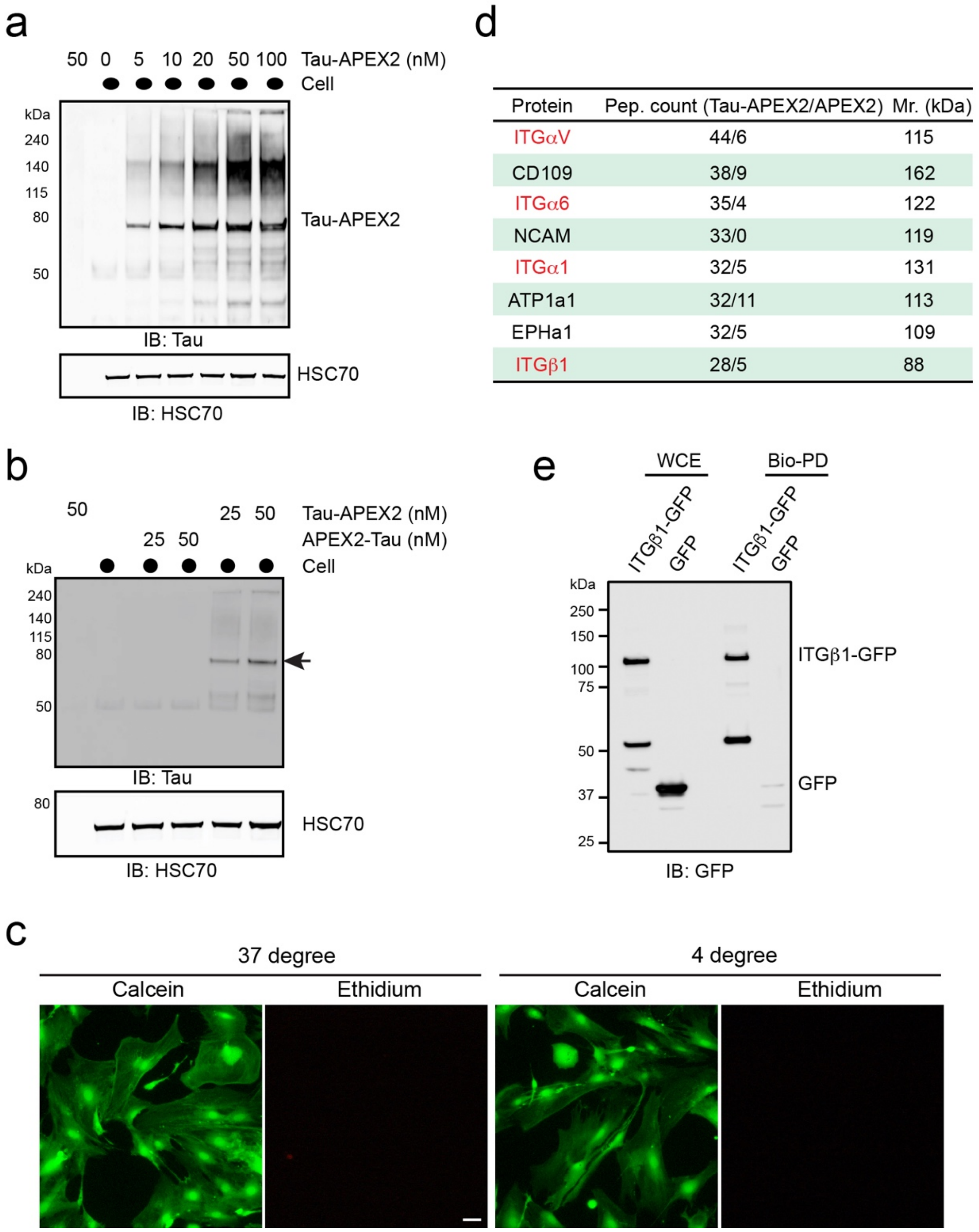


| Protein | Pep. count (Tau-APEX2/APEX2) | Mr. (kDa) |
|---|---|---|
| ITGαV | 44/6 | 115 |
| CD109 | 38/9 | 162 |
| ITGα6 | 35/4 | 122 |
| NCAM | 33/0 | 119 |
| ITGα1 | 32/5 | 131 |
| ATP1a1 | 32/11 | 113 |
| EPHa1 | 32/5 | 109 |
| ITGβ1 | 28/5 | 88 |

**Figure S2. Identification of integrin αV/β1 as a Tau interactor in primary astrocytes.**

(**a**) Tau-APEX2 binds to the cell surface in a dose dependent manner. Sh-SY5Y cells were incubated with the indicated amount of Tau-APEX2 on ice for 3 h. After washing, cells were lysed and cell lysates were analyzed by immunoblotting. As a negative control, 50 nM Tau PFF was incubated with buffer (no cells).

(**b**) Tau-APEX2 but not APEX2-Tau binds to the cell surface. Sh-SY5Y cells were incubated with the indicated amount of Tau protein on ice for 2 h, washed, and then lysed. Cell lysates were analyzed by immunoblotting. The arrow indicates Tau-APEX2.

(**c**) Short-term treatment of astrocytes with low temperature did not induce cell death. Primary astrocytes treated at 37 °C or 4 °C for 1 h were stained with calcein-AM and ethidium homodimer-1. Scale bar, 10 µm.

(**d**) A summary of the top candidates identified by mass spectrometry.

(**e**) Validation of Tau-APEX2-mediated biotinylation of ITGβ1. Tau-APEX2 was incubated with HEK293T cells transfected with either an empty vector (EV) or an ITGβ1-GFP-expressing plasmid followed by *in vitro* biotinylation. Whole cell extracts (WCE) were either directly analyzed by immunoblotting or first subjected to biotin pulldown (PD) before immunoblotting with GFP antibodies.

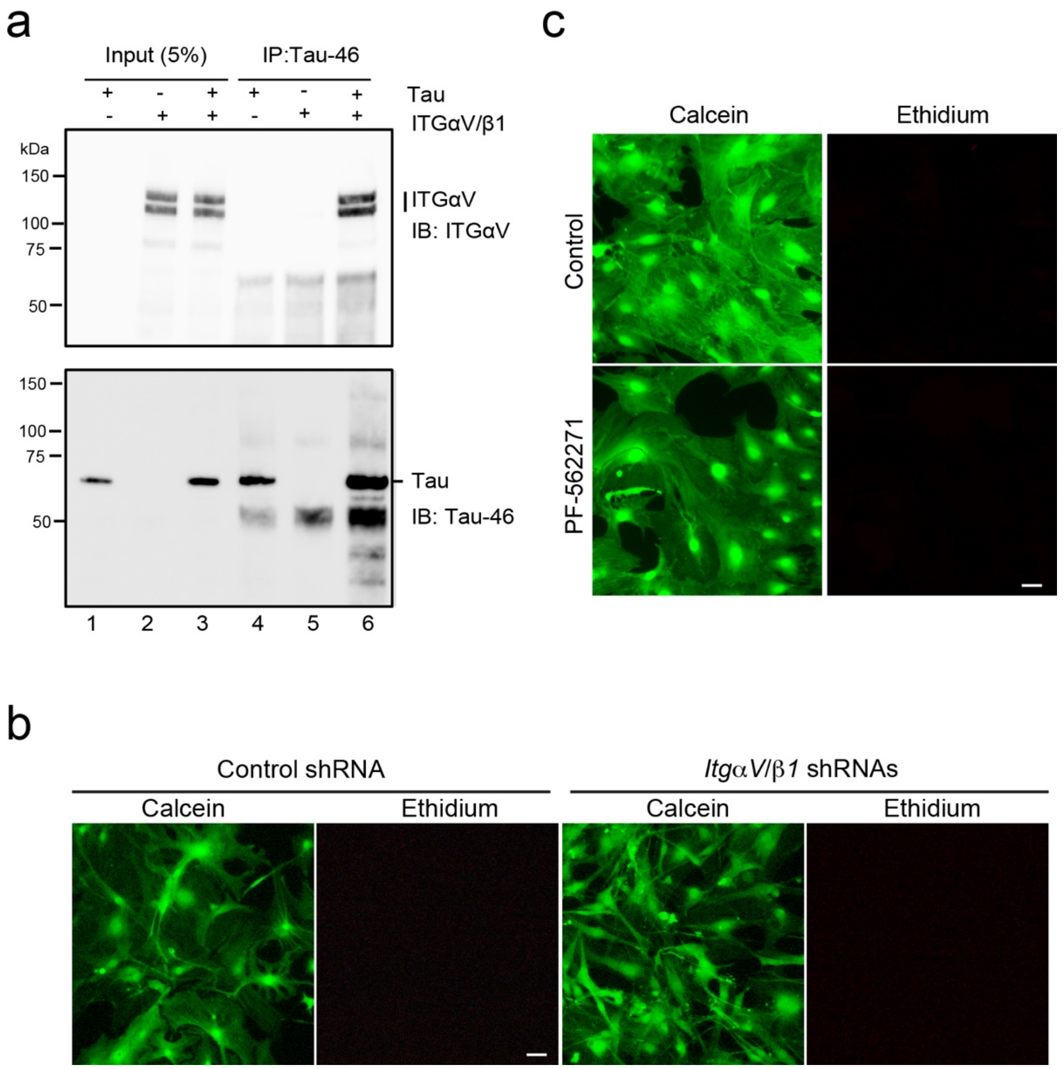


**Figure S3. Tau binds integrin αV/β1 directly in vitro.**

(**a**) Recombinant integrin αVβ1 (100 nM) was incubated with PBS as a control or 100 nM monomeric recombinant Tau protein as indicated. A fraction of the binding reaction (5% input) was analyzed directly by immunoblotting (lanes 1-3). The remaining samples were subjected to immunoprecipitation with Tau-46 antibodies before immunoblotting (lanes 4-6).

(**b**) The effect of integrin αV/β1 knockdown on astrocyte viability. Cells infected with lentiviruses expressing either control or integrin αV/β1 shRNAs for 72 h were stained with calcein-AM and ethidium homodimer-1. Scale bar, 10 μm.

(**c**) The effect of FAK inhibition on astrocyte viability. Cells treated with the indicated drugs  for 8 h were stained with calcein-AM and ethidium homodimer-1. Scale bar, 10 μm.

a

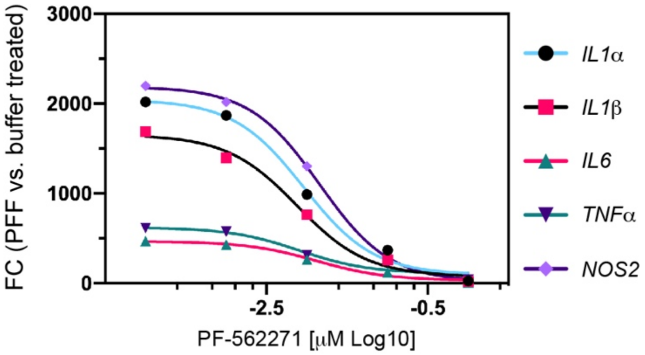


b

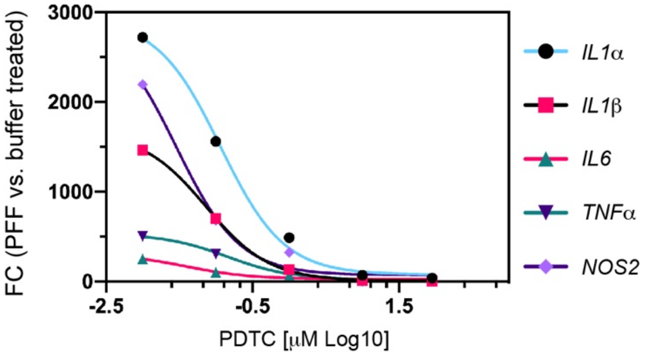

**Figure S4. Dose dependent suppression of Tau-PFF-induced inflammation by PF-562271 and PDTC.**

Immunopurified astrocytes were pre-treated with the indicated concentration of PF-562271 (**a**) or PDTC (**b**) for 1h and then treated with either PBS or Tau PFF. The expression of the indicated genes was determined by qRT-PCR and normalized to PBS-treated samples.

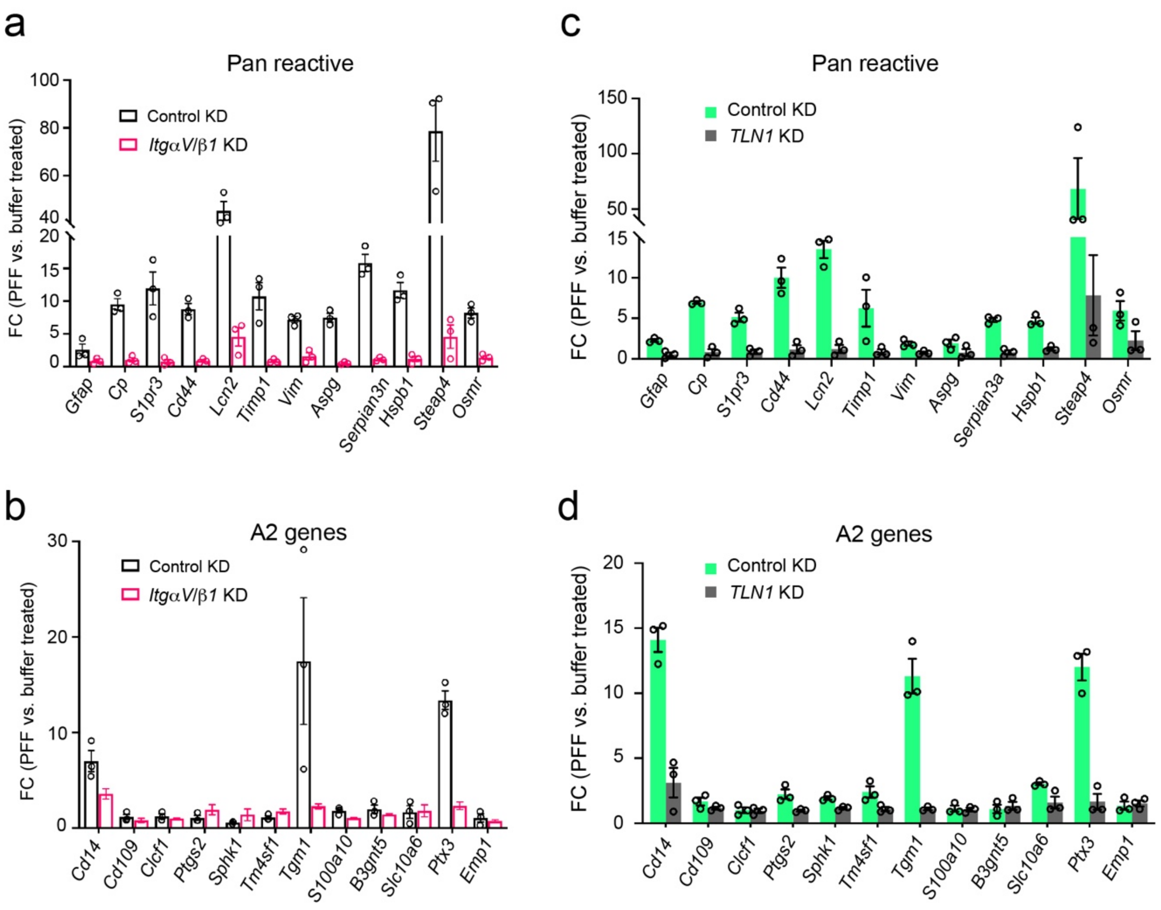


**Figure S5. The effect of integrin αV/β1 and Talin1 knockdown on the expression of pan-reactive and A2 specific genes in astrocytes.**

(**a, b**) Control or integrin αV/β1 knockdown astrocytes were treated with either PBS or Tau PFFs. The expression of the indicated genes was determined by qRT-PCR and normalized to PBS-treated samples. Mean ±SEM, n=3 independent experiments.

(**c**, **d**) Control or Talin1(TLN) knockdown immunopurified astrocytes were treated with either PBS or Tau PFFs. The expression of the indicated genes was determined by qRT-PCR and normalized to PBS-treated samples. Mean ±SEM, n=3 independent experiments.

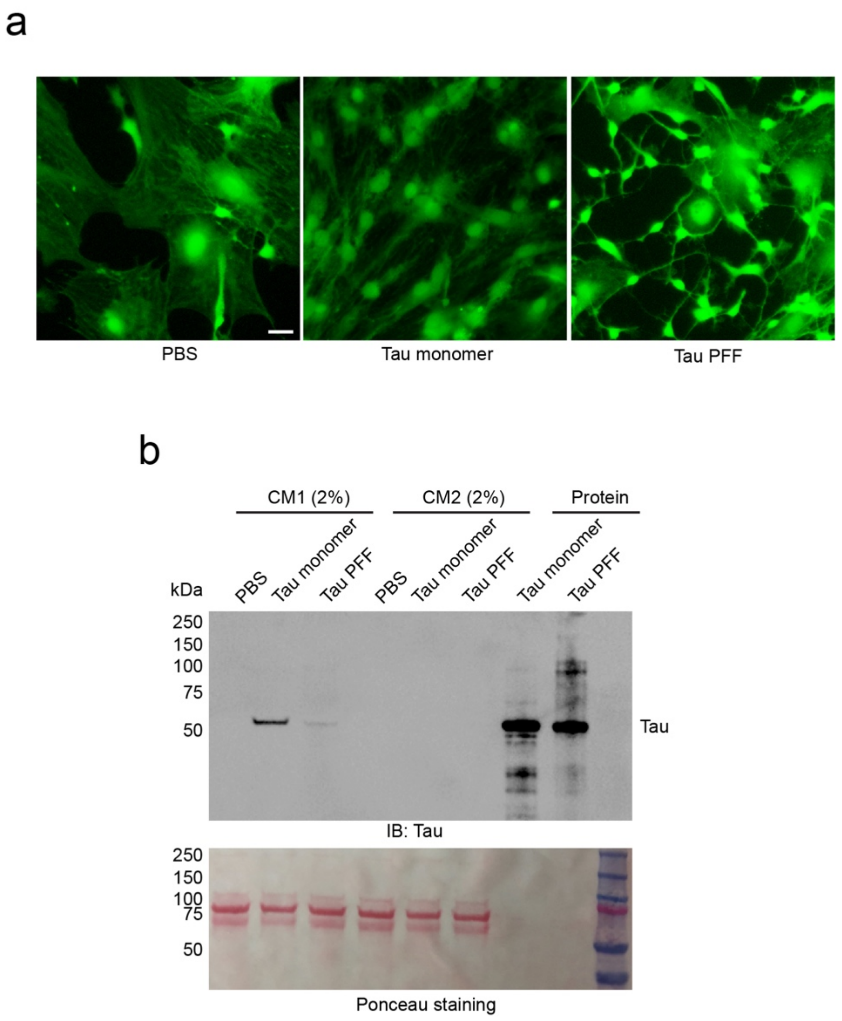

**Figure S6 Tau PFF-treated astrocytes release a neurotoxic factor(s).**

(**a**) Morphological changes in Tau PFF-treated astrocytes. Primary astrocytes were treated with PBS, Tau monomer or PFF (200nM, 6 h) and then replated in a new dish with fresh medium for 48 h before calcein-AM staining. Scale bar, 10 μm.

(**b**) No Tau carryover was detected in conditioned medium (CM) from Tau-treated astrocytes. Astrocytes were treated with PBS, monomeric Tau (200nM) or Tau PFF (200nM) for 6h. Conditioned medium (CM1) was harvested. Cells were then washed, trypsinized, and transferred to a new plate. After cell attachment, conditioned medium (CM2) was harvested. The CM1 and CM2 were analyzed by immunoblotting together with purified Tau monomer and PFF as a control.

**SI Table 1 A list of reagents**

| REAGENTS | SOURCE |
|---|---|
| **PLASMIDS** | |
| pcDNA3 APEX2-NES | A gift from Alice Ting Addgene (# 49386) [1] |
| TAU/PET29B | A GIFT FROM PETER KLEIN ADDGENE (# 16316) [2] |
| pEF1-αV | A gift from Timothy Springer Addgene (# 27290) [3] |
| Integrin-β1-GFP | A gift from Martin Humphries Addgene (# 69804) [4] |
| pCMV-VSV-G | A gift from Bob Weinberg Addgene (#8454) [5] |
| pSPAX2 | A gift from Didier Trono Addgene (#12260) |
| pEGFP-C1 | Clonetech |
| ITGαV MISSION SHRNA SHRNA PLASMID DNA | Sigma SHCLND-NM_008402 TRCN0000066589 |
| ITGβ1 MISSION SHRNA SHRNA PLASMID DNA | Sigma SHCLND-NM_010578 TRCN0000348624 |
| TLN1 MISSION SHRNA SHRNA PLASMID DNA | Sigma SHCLND-NM_011602 TRCN0000108756 |
| **CHEMICALS** | |
| PUROMYCIN | Sigma |
| PDTC | R&D |
| PH RODO-RED SUCCINIMIDYL ESTER | ThermoFisher Scientific |
| ALEXA 596 SUCCINIMIDYL ESTER | ThermoFisher Scientific |
| DYNASORE | TOCRIS |
| JASPLAKINOLIDE | TOCRIS |
| HOECHEST 33342 | ThermoFisher Scientific |
| IPTG | TOCRIS |
| PF-562271 | Selleckchem |
| BIOTIN-PHENOL | Iris-Biotech Cat # LS-3500.0250 |
| **ANTIBODIES (Dilution)** | |
| INTEGRIN αV (1:1,000) | Abcam Cat # ab179475 |
| INTEGRIN β1 (1:1,000) | Abcam Cat # ab52971 |
| NFκB P65 (1:500) | Rockland Cat# 200-301-065 |
| INTEGRIN β5 (20 μg/immune-panning) | R&D Systen Cat# AF3824 |

| | |
|---|---|
| GFAP (1:250) | Proteintech Cat # 60190 |
| TAU (TAU 46) (1:1,000) | Santa Cruz Cat # sc-32274 |
| TAU (TAU H150) (2 μg/IP) | Santa Cruz Cat # sc-5587 |
| BIOTIN (BTN,4) (1:1,000) | Thermo Fisher Cat # MA5-11251 |
| FLAG (M2) (1:1,000) | Sigma Cat # F1804-200UG |
| GFP (B2) (1:500) | Santa Cruz Cat# SC-9996 |
| HSP90 (1:1,000) | Santa Cruz Cat # sc-69703 |
| CD45 (1.25 μg/immune-panning) | BD Biosciences Cat# 550539 |
| ANTI-MOUSE IgG PEROXIDASE ANTIBODY (1:5,000) | Sigma Cat# A4416-1ML |
| ANTI-RABBIT IgG PEROXIDASE ANTIBODY (1:5,000) | Sigma Cat# A6154-1ML |
| GOAT ANTI-MOUSE IgG (H+L) ALEXA $FLUOR_{680}$ (1:10,000) | Thermo Fisher Cat# A21058 |
| GOAT ANTI-RABBIT IgG (H&L) $DYLIGHT_{800}$ CONJUGATED (1:10,000) | Rockland Cat# 611-145-122 |
| GOAT ANTI-MOUSE IgG ALEXA $FLUOR_{488}$ (1:1,000) | Thermo Fisher Cat# A21121 |
| GOAT ANTI-RAT IgG (H + L) (80 μg/ immune-panning) | Jackson ImmunoResearch Cat# 112-005-167 |
| DONKEY ANTI-SHEEP IGG (H + L) (80 μg/ immune-panning) | Jackson ImmunoResearch Cat# 713-005-147 |
| **REGENTS** | |
| STREPTAVIDIN MAGNETIC BEADS | Pierce Cat # 88817 |
| SODIUM ASCORBATE | VWR International Cat # 95035-692 |
| TROLOX | Sigma-Aldrich Cat. # 238813-5G |
| SODIUM AZIDE | VWR International Cat # AA14314-22 |
| HYDROGEN PEROXIDE 30% (WT/WT) | Sigma-Aldrich Cat # H1009-100ML |
| **PROTEIN** | |
| RECOMBINANT HUMAN INTEGRIN ALPHA 1 BETA 1 PROTEIN | R&D system |

**SI Table 2. qPCR primers used in the study**

| | |
|---|---|
| IL1α-F | 5’-GGGAAGATTCTGAAGAAGAG |
| IL1α-R | 5’-GAGTAACAGGATATTTAGAGTCG |
| IL1β-F | 5’-TGTGAAATGCCACCTTTTGA |
| IL1β-R | 5’-GTGCTCATGTCCTCATCCTG |
| IL6-F | 5’-GACAACCACGGCCTTCCCTACTTC |
| IL6-R | 5’-TCATTTCCACGATTTCCCAGAGA |
| TNFα-F | 5’-CCGATGGGTTGTACCTTGTC |
| TNFα-R | 5’-CGGACTCCGCAAAGTCTAAG |
| IL10-F | 5’-AAGGCAGTGGAGCAGGTGAA |
| IL10-R | 5’-CCAGCAGACTCAATACACAC |
| NOS2-F | 5’-CACCTTGGAGTTCACCCAG |
| NOS2-R | 5’-ACCACTCGTACTTGGGATGC |
| TGFβ1-F | 5’-TACCATGCCAACTTCTGTCTGGGA |
| TGFβ1-R | 5’-TGTGTTGGTTGTAGAGGGCAAGGA |
| CCL2-F | 5’-TCAGCCAGATGCAGTTAACG |
| CCL2-R | 5’-GATCCTCTTGTAGCTCTCCAGC |
| CCL3-F | 5’-GACTGCCTGCTGCTTCT |
| CCL3-R | 5’-GATCTGCCGGTTTCTCTTAG |
| CCL4-F | 5’-CATGAAGCTCTGCGTGTCT |
| CCL4-R | 5’-CTGCCGGGAGGTGTAA |
| CXCL10-F | 5’-GCCGTCATTTTCTGCCTCAT |
| CXCL10-R | 5’-GCTTCCCTATGGCCCTCATT |
| CCL12-F | 5’-AGAATCACAAGCAGCCAGTGT |

| | |
|---|---|
| CCL12-R | 5'-ATCCAAGTGGTTTATGGAATTCTTAAC |
| ITGβ1-F | 5'-ATGCCAAATCTTGCGGAGAAT |
| ITGβ1-R | 5'-TTTGCTGCGATTGGTGACATT |
| ITGαV-F | 5'-CCTGAGACTGAAGAAGAC |
| ITGαV-R | 5'-CCTTGCTGAATGAACTTG |
| ITGβ5-F | 5'-TGACGAAGAACCACTATA |
| ITGβ5-R | 5'-CTACTGTACGCATTGATAA |
| ITGα6-F | 5'-AAGGAAGGATGTGGAGAC |
| ITGα6-R | 5'-TTGAATTGGAAGGTAAGAGAAT |
| ITGα1-F | 5'-AGCCTATCCTGAGACCTT |
| ITGα1-R | 5'-TCTTATCTTCACCACAGTTCT |
| ITGα3-F | 5'-GAGCTGTGGTTGGTGCTTG |
| ITGα3-R | 5'-GCACTTCCACAAGAGGAGGAT |
| TLN1-F | 5'-CTGGCCTCACAAGCCAAG |
| TLN1-R | 5'-TTGATGTGAGCGCCTATCTCT |
| Iigp1-F | 5'-GGGGCAATAGCTCATTGGTA |
| Iigp1-R | 5'-ACCTCGAAGACATCCCCTTT |
| Gbp2-F | 5'-GGGGTCACTGTCTGACCACT |
| Gbp2-R | 5'-GGGAAACCTGGGATGAGATT |
| Fbln5-F | 5'-CTTCAGATGCAAGCAACAA |
| Fbln5-R | 5'-AGGCAGTGTCAGAGGCCTTA |
| Ugt1a-F | 5'-CCTATGGGTCACTTGCCACT |
| Ugt1a-R | 5'-AAAACCATGTTGGGCATGAT |
| Psmb8-F | 5'-CAGTCCTGAAGAGGCCTACG |
| Psmb8-R | 5'-CACTTTCACCCAACCGTCTT |

| | |
|---|---|
| Srgn-F | 5’-GCAAGGTTATCCTGCTCGGA |
| Srgn-R | 5’-TGGGAGGGCCGATGTTATTG |
| Amigo2-F | 5’-GAGGCGACCATAATGTCGTT |
| Amigo2-R | 5’-GCATCCAACAGTCCGATTCT |
| Clcf1-F | 5’-CTTCAATCCTCCTCGACTGG |
| Clcf1-R | 5’-TACGTCGGAGTTCAGCTGTG |
| Tgm1-F | 5’-CTGTTGGTCCCGTCCCAAA |
| Tgm1-R | 5’-GGACCTTCCATTGTGCCTGG |
| S100A10-F | 5’-CCTCTGGCTGTGGACAAAAT |
| S100A10-R | 5’-CTGCTCACAAGAAGCAGTGG |
| Sphk1-F | 5’-GATGCATGAGGTGGTGAATG |
| Sphk1-R | 5’-TGCTCGTACCCAGCATAGTG |
| Slc10a6-F | 5’-GCTTCGGTGGTATGATGCTT |
| Slc10a6-R | 5’-CCACAGGCTTTTCTGGTGAT |
| Tm4sf1-F | 5’-GCCCAAGCATATTGTGGAGT |
| Tm4sf1-R | 5’-AGGGTAGGATGTGGCACAAG |
| B3gnt5-F | 5’-CGTGGGGCAATGAGAACTAT |
| B3gnt5-R | 5’-CCCAGCTGAACTGAAGAAGG |
| VCAM-F | 5’-TCTGGGAAGCTGGAACGAAG |
| VCAM-R | 5’-CAAACACTTGACCGTGACCG |
| ITGαM-F | 5’-ATGGACGCTGATGGCAATACC |
| ITGαM-R | 5’-TCCCCATTCACGTCTCCCA |
| Serping1-F | 5’-ACAGCCCCCTCTGAATTCTT |
| Serping1-R | 5’-GGATGCTCTCCAAGTTGCTC |
| NF-κB-F | 5’-CCAACGCCCTCTTCGACTAC |

| | |
|---|---|
| NF-κB-R | 5’-GATCCCTCACGAGCTGAGC |
| GFAP-F | 5’-AGAAAGGTTGAATCGCTGGA |
| GFAP-R | 5’-CGGCGATAGTCGTTAGCTTC |
| CD109-F | 5’-CACAGTCGGGAGCCCTAAAG |
| CD109-R | 5’-GCAGCGATTTCGATGTCCAC |
| CD14-F | 5’-GGACTGATCTCAGCCCTCTG |
| CD14-R | 5’-GCTTCAGCCCAGTGAAAGAC |
| H2-D1-F | 5’-TCCGAGATTGTAAAGCGTGAAGA |
| H2-D1-R | 5’-ACAGGGCAGTGCAGGGATAG |
| H2-T23-F | 5’-GGACCGCGAATGACATAGC |
| H2-T23-R | 5’-GCACCTCAGGGTGACTTCAT |
| Emp1-F | 5’-GAGACACTGGCCAGAAAAGC |
| Emp1-R | 5’-TAAAAGGCAAGGGAATGCAC |
| Fkbp5-F | 5’-TATGCTTATGGCTCGGCTGG |
| Fkbp5-R | 5’-CAGCCTTCCAGGTGGACTTT |
| CD44-F | 5’-TCAGGATAGCCCCACAACAAC |
| CD44-R | 5’-GACTCCGTACCAGGCATCTTC |
| Serpina3n-F | 5’-GTCTTTCAGGTGGTCCACAAGG |
| Serpina3n-R | 5’-GCCAATCACAGCATAGAAGCG |
| CP-F | 5’-GATGTTTCCCCAAACGCCTG |
| CP-R | 5’-GTAGCTCTGAGACGATGCTTGA |
| S1pr3-F | 5’-CTTGCAGAACGAGAGCCTGT |
| S1pr3-R | 5’-CCTCAACAGTCCACGAGAGG |
| GFAP-F | 5’-AACCGCATCACCATTCCTGT |
| GFAP-R | 5’-TCCTTAATGACCTCGCCATCC |

| | |
|---|---|
| Lcn2-F | 5’-CCGACACTGACTACGACCAG |
| Lcn2-R | 5’-AATGCATTGGTCGGTGGGAA |
| Timp1-F | 5’-CGCTAGAGCAGATACCACGA |
| Timp1-R | 5’-CCAGGTCCGAGTTGCAGAAA |
| Vim-F | 5’-GAGGAGATGAGGGAGTTGCG |
| Vim-R | 5’-CTGCAATTTTTCTCGCAGCC |
| ASPG-F | 5’-CAGGTGCCCAGGTTCCTATC |
| ASPG-R | 5’-GTCCACCTTGGTTGTCCGAT |
| Hsbp1-F | 5’-GAGATCACTGGCAAGCACGA |
| Hsbp1-R | 5’-ATTGTGTGACTGCTTTGGGC |
| Steap4-F | 5’-CAAACGCCGAGTACCTTGCT |
| Steap4-R | 5’-CAGACAAACACCTGCCGACT |
| Osmr-F | 5’-GTCATTCTGGACATGAAGAGGT |
| Osmr-R | 5’-AATCACAGCGTTGGGTCTGA |
